# Tracing cell communication programs across conditions at single cell resolution with CCC-RISE

**DOI:** 10.64898/2026.04.14.718551

**Authors:** Andrew Ramirez, Nathaniel Thomas, Daniel R. Calabrese, John R. Greenland, Aaron S. Meyer

## Abstract

Cell-cell communication (CCC) mediates coordinated cellular activities that vary dynamically across time, location, and biological context. While various tools exist to infer CCC, they typically aggregate data according to pre-defined cell types, obscuring critical single-cell heterogeneity. Furthermore, because signaling pathways and cell populations operate in a coordinated manner, an integrative analytical approach is essential. To address these challenges, we developed CCC-RISE, an extension of the tensor-based method Reduction and Insight in Single-cell Exploration (RISE). CCC-RISE identifies integrative patterns of single-cell variation by deconvolving communication into interpretable modules defined by unique sender cells, receiver cells, ligands, and condition associations. We applied this framework to a COVID-19 cohort with varying disease severity and a lung transplant cohort with acute allograft dysfunction. In both contexts, CCC-RISE successfully identified disease-relevant communication programs and traced them to specific cellular subpopulations, often crossing conventional cell-type boundaries. This approach offers a robust pipeline enabling the identification of disease-relevant signaling subpopulations that are invisible to aggregate methods.

**Highlights:**

- CCC-RISE enables integrative analysis of cell-cell communication across multiple conditions at single-cell resolution
- CCC-RISE deconvolves signaling patterns into modules defined by their sender cells, receiver cells, LR pairs, and experimental conditions/samples
- Analysis at single-cell resolution uncovers signaling activity within and across conventional cell types

## INTRODUCTION

Multicellular organisms rely on coordinated cell-cell communication (CCC) to guide development, maintain homeostasis, and mount immune responses^1^. This communication is mediated by a diverse molecular toolkit, including ligands, receptors, chemokines, hormones, and structural proteins^1,2^. CCC is initiated when extracellular signals trigger intracellular cascades, often altering gene expression to regulate cellular behavior and function. By mapping how diseases disrupt these pathways, we can characterize the specific cellular responses that drive and define pathological states.

To better understand how cells interact, researchers have developed various computational techniques to infer cell-cell communication (CCC) from transcriptomic data^3,4^. These methods typically leverage single-cell RNA sequencing (scRNA-seq) to identify the co-expression of ligand-receptor (LR) gene pairs documented in curated databases^5,6^. This approach hinges on the central assumption that transcript levels accurately reflect functional protein abundance, which in turn determines the probability of interaction. While this is a significant simplification of biological complexity, it allows for the high-throughput screening of thousands of interactions, providing a data-driven way to prioritize specific communication events for experimental validation^7,8^. Initial tools such as CellPhoneDB and CellChat use statistical tests to compare communication scores against a null distribution created by randomly permuting cell labels^5,9^. Originally designed to detect the presence of signaling pathways, these foundational tools have since inspired a diverse analytical toolkit capable of addressing increasingly complex biological questions^10^.

A critical objective of CCC analysis is to understand how cellular interactions evolve across distinct biological contexts, such as disease states, developmental stages, or treatment groups^11–14^. By mapping alterations in ligand-receptor activity and signaling pathways, researchers can pinpoint the communication programs that drive phenotypic outcomes and identify potential therapeutic targets. Computational approaches for multi-condition CCC analysis generally fall into two primary categories: matrix-based and tensor-based methods. Matrix-based strategies—including DIALOGUE, MultiNicheNet, and Multicellular Factor Analysis—typically flatten complex data into two-dimensional arrays, such as rows of samples against columns of sender-receiver features^15–17^. While these methods can successfully identify statistically significant ligand-receptor pairs, the flattening process creates a confounding mixture of signals that makes it difficult to track context-dependent communication events across specific sender and receiver cell types. In contrast, tensor-based methods like Tensor-cell2cell, scTensor, STACCato, and scITD address this limitation by modeling data across all dimensions simultaneously. By arranging data into higher-order tensors rather than flat matrices, these methods preserve the multi-dimensional structure of the interaction, allowing for the identification of coordinated changes in CCC and the regulation of communication programs across conditions^11,18–20^. To extract these insights, tensor-based tools employ factorization techniques, such as Canonical Polyadic Decomposition (CPD), which model the tensor as a sum of rank-one components to reveal distinct signaling patterns^21,22^.

Despite the advantages of current tensor-based methods, their reliance on pseudobulking single-cell data represents a significant bottleneck. Because standard decompositions like CPD require data dimensions to be strictly aligned (i.e., “sender cell type 1” must match perfectly across all conditions) researchers are forced to aggregate raw single-cell data into discrete categories. This approach relies on the reductive assumption that a cell can be fully defined by a single label^8^. In biological reality, cells are multifaceted entities that often hold simultaneous identities, blending stable lineage markers with transient functional states driven by their immediate environment^23–25^. Furthermore, the process of annotation itself is fraught with challenges; cell type labeling is inevitably incomplete or inaccurate, particularly when researchers attempt to define fine-grained labels to capture subtle heterogeneity. By forcing cells into these rigid, pre-defined boxes, pseudobulking obscures the functional diversity essential for understanding cellular communication, as distinct subpopulations within a single “type” may participate in entirely different signaling programs. Conversely, while methods such as Scriabin, DeepCOLOR, and NICHES allow for analysis at single-cell resolution, they lack the ability to model how communication programs are regulated across biological contexts, a key strength of tensor-based approaches^26–30^. Therefore, there is a pressing need for a framework that can interpret the multi-dimensional nature of CCC across conditions while preserving the resolution necessary to capture the complex, overlapping identities of individual cells.

To address these limitations, here we developed CCC-RISE, a framework designed to bridge the gap between multi-condition analysis and single-cell resolution. CCC-RISE builds upon the Reduction and Insight in Single-cell Exploration (RISE) method, which utilizes PARAFAC2 to analyze scRNA-seq data without requiring predefined cell type labels or aggregation^31^. PARAFAC2 aligns cells across conditions by solving an Orthogonal Procrustes problem, projecting each cell onto a shared set of latent expression programs, termed “eigen-states”, that capture the dominant axes of transcriptional variation common to all samples. Each eigen-state represents a weighted combination of genes, and each cell receives a loading across all eigen-states, preserving continuous variation rather than forcing discrete assignments. By applying CCC scoring to these eigen-states, we can identify core communication programs across varying conditions while preserving the ability to interpret these signals at the individual cell level. We demonstrate the utility of CCC-RISE in two clinical applications, effectively linking coordinated communication signatures to relevant patient features and offering a precise new approach to dissecting CCC dynamics across biological contexts.

## RESULTS

### CCC-RISE framework overview and validation strategy

RISE projects cells from multiple conditions onto a common reference frame, effectively aligning them based on gene expression without requiring *a priori* labeling. To overcome the limitations of aggregating data into pre-defined cell types, CCC-RISE builds upon the unsupervised “eigen-states” generated by the RISE framework (Fig. 1a)^31^. CCC-RISE computes communication scores directly on these intermediate eigen-states rather than on individual cells or broad populations. Because RISE projections summarize the common gene expression profiles across the eigen-states, these matrices can serve as the basis for an overall cell embedding (Fig. 1b). For every experimental condition and ligand-receptor (LR) pair, a score is calculated for all combinations of sender and receiver eigen-states by integrating the ligand expression of the sender with the receptor expression of the receiver. This process constructs a four-dimensional tensor describing the data across sender eigen-states, receiver eigen-states, experimental conditions, and LR pairs.

**Figure 1.**
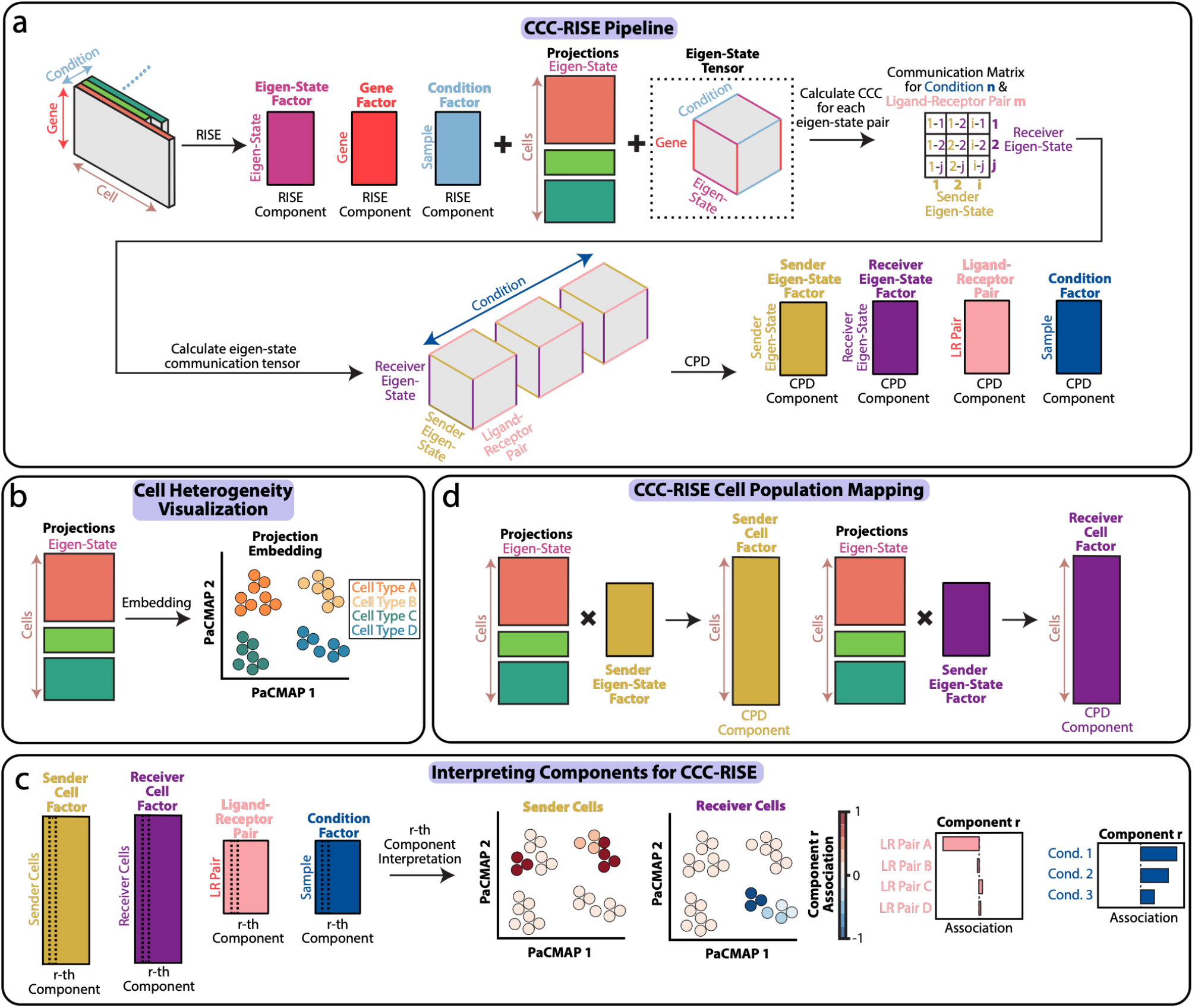
CCC-RISE deconvolves cell communication variability across single cells and conditions. **(a)** The CCC-RISE workflow. The RISE method first projects multi-condition scRNA-seq data into eigen-states to model cell heterogeneity. Communication scores are then computed between these eigen-states for ligand-receptor (LR) pairs to form a 4D tensor. Finally, CPD factorization decomposes this tensor into distinct factors that describe the component associations with sender eigen-states, receiver eigen-states, LR pairs, and conditions. **(b)** RISE projections serve as a common embedding basis to visualize cell-to-cell heterogeneity across all conditions. **(c)** Interpreting communication programs. Each extracted component is defined by the association weights across four dimensions: sender cells, receiver cells, LR pairs, and experimental conditions. **(d)** Linking factors to cells. Sender and receiver eigen-state factors are mathematically combined with the RISE projections to map communication patterns back to specific single-cell populations.

To extract patterns from this high-dimensional structure, the method employs Canonical Polyadic Decomposition (CPD), which factors the 4D tensor into distinct components. Similar to how principal component analysis (PCA) reduces dimensionality, this decomposition approximates the tensor as a sum of rank-one tensors, each representing a coordinated communication program defined by specific variations in sender and receiver states, LR pairs, and conditions (Fig. 1c)^21,22,32^. Crucially, because the original RISE projections preserve cell-to-cell heterogeneity, the component loadings on eigen-states can be projected back onto individual cells. By multiplying the RISE projections with the sender and receiver eigen-state factors, the specific association of individual cells with each communication component is recovered. This allows for the identification of specific cell subpopulations driving communication signals, enabling the tracing of signaling programs across conditions without losing the granularity of single-cell data (Fig. 1d).

### CCC-RISE Uncovers Subpopulation-Specific Immune Signaling Heterogeneity in COVID-19

To evaluate the ability of CCC-RISE to resolve communication patterns at single-cell resolution, we applied the framework to twelve bronchoalveolar lavage (BAL) samples collected from patients with varying degrees of COVID-19 severity (healthy, moderate, and severe; Fig. 2a)^33^. This dataset was previously used for equivalent analysis in pseudobulk form^11^. These samples were characterized by distinct cellular compositions, notably a significant increase in epithelial cell proportions and a decrease in myeloid-derived suppressor cells (mDCs) corresponding with disease severity (Supplementary Fig. 1a-c). We hypothesized that CCC-RISE would not only capture these broad shifts but also uncover high-resolution communication patterns driven by specific sender and receiver subpopulations that might be obscured within conventional cell type annotations.

**Figure 2.**
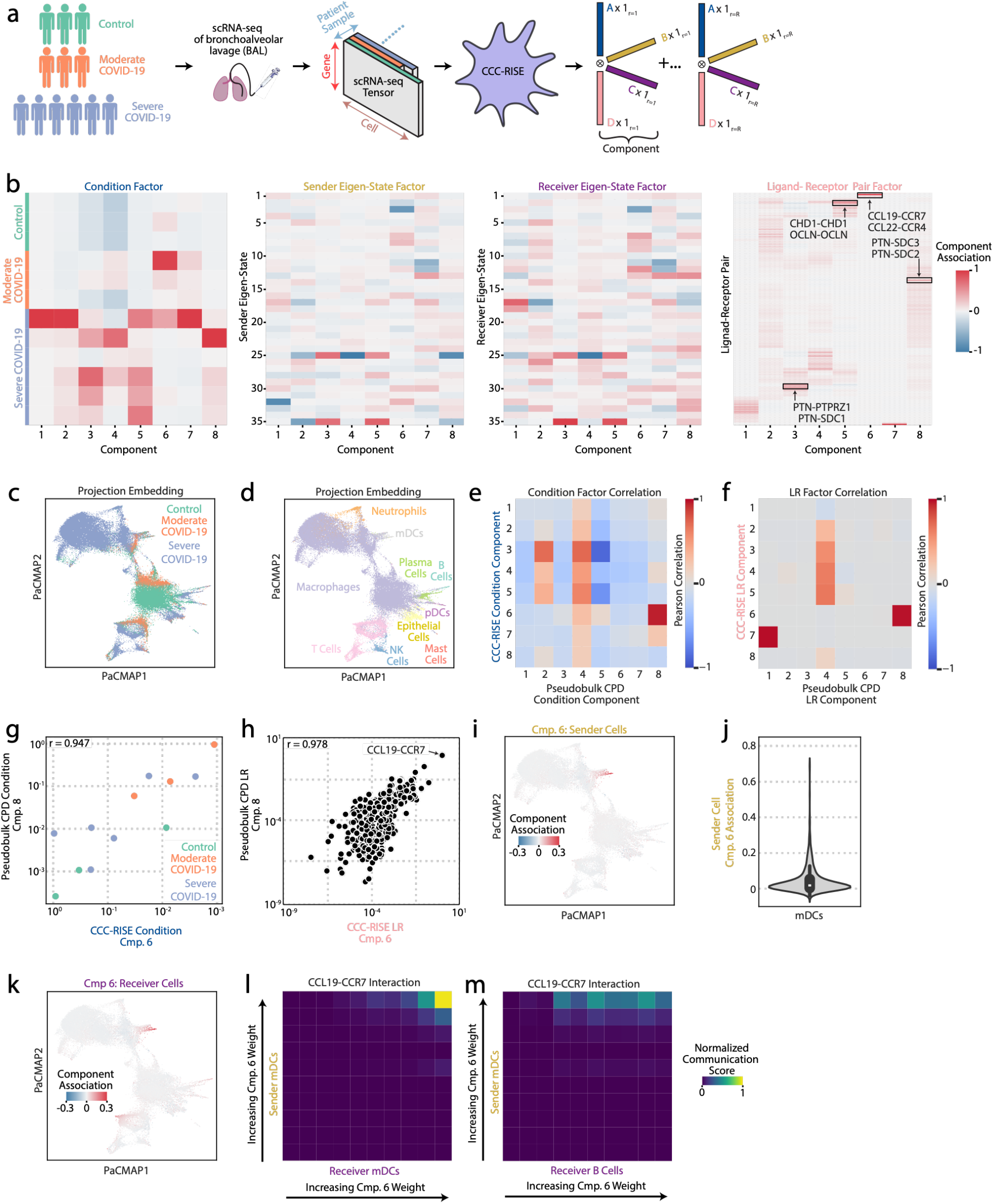
CCC-RISE resolves COVID-19-associated communication patterns to specific immune subpopulations. **(a)** Schematic of the CCC-RISE workflow applied to bronchoalveolar lavage (BAL) samples from patients with healthy, moderate, and severe COVID-19. **(b)** Decomposed factor matrices representing the conditions, sender eigen-states, receiver eigen-states, and ligand-receptor (LR) pairs for each identified component. **(c–d)** PaCMAP embeddings of the RISE projections, visualizing cell-to-cell heterogeneity colored by (c) patient severity group and (d) cell-type annotations. **(e-f)** Heatmaps displaying the Pearson correlation between components identified by CCC-RISE and those from pseudobulk CPD analysis for (e) condition factors and (f) LR pair factors. **(g)** Correlation between CCC-RISE Component 6 and Pseudobulk Component 8 condition factors, annotated with Pearson correlation coefficients and colored by patient group. **(h)** Correlation between CCC-RISE Component 6 and Pseudobulk Component 8 LR pair factors, highlighting the top-weighted pair (CCL19-CCR7). **(i)** PaCMAP embedding of RISE projections with sender cells colored by their specific loadings in Component 6. **(j)** Distribution of Component 6 loadings specifically within the sender mDC population, showing intra-type heterogeneity. **(k)** PaCMAP embedding of RISE projections with receiver cells colored by their loadings in Component 6. **(l-m)** Normalized communication scores for the CCL19-CCR7 interaction in Component 6, visualized for (l) sender mDCs to receiver mDCs, and (m) sender mDCs to receiver B cells.

To ensure the robustness of the decomposition, we first optimized the model parameters using a bootstrapping strategy to measure the stability of the factor matrices. We selected 35 RISE components to capture sufficient variance while maintaining factor stability, and subsequently decomposed the resulting communication tensor into an 8-component CPD model, which explained over 25% of the variance with high reproducibility (Supplementary Fig. 1d-g). This process yielded distinct factor matrices for conditions, sender eigen-states, receiver eigen-states, and LR pairs (Fig. 2b), with each component delineating a specific cell communication program (Supplementary Figs. 2 and 3). To visualize the cellular heterogeneity underlying these components, we generated embeddings based on the RISE projections, which effectively clustered cells into distinct immune lineages (Fig. 2c-d).

We next benchmarked the resolution of CCC-RISE against the pseudobulk tensor decomposition equivalent. In the pseudobulk approach, scRNA-seq data was aggregated by cell type before LR scoring and factorization (Supplementary Fig. 1h-i). While we expected both methods to identify core signaling programs, we anticipated that CCC-RISE would provide superior granularity. Indeed, we observed a strong correlation between CCC-RISE Component 6 and Pseudobulk Component 8, indicating that both methods captured the same underlying condition-specific pattern (Fig. 2e-h). Both approaches highlighted mDCs as key sender cells and both mDCs and B cells as primary receivers (Supplementary Fig. 1i). However, the pseudobulk analysis uniformly implicated the entire annotated cell populations.

In contrast, CCC-RISE revealed significant heterogeneity within these canonical cell types. By mapping the component loadings back to individual cells, we observed that only a specific subset of the annotated mDC and B cell populations was driving this communication signal (Fig. 2i-j). This was further evidenced by the non-uniform distribution of communication scores for the top-weighted LR pair, CCL19-CCR7, which were concentrated within these active subpopulations rather than being present across the entire cell type (Fig. 2k-m). This interaction is consistent with the established role of mDCs in recruiting lymphocytes to initiate adaptive immunity, a process known to be critical in viral pathogenesis^34–37^. Furthermore, the absence of CCL19-CCR7 interactions in non-relevant populations, such as macrophages and NK cells, confirmed that the CCC-RISE model correctly avoided an association with these cell types (Supplementary Fig. 1j–k).

### CCC-RISE reveals severity-associated communication programs in COVID-19 at single-cell resolution

Following the comparison against pseudobulk analysis, we sought to identify specific communication programs tracking with disease progression. We found that condition factors for Components 3, 5, and 8 exhibited the strongest correlation with COVID-19 severity (Fig. 3a). We first examined Component 3, which was predominantly driven by epithelial cells acting as both senders and receivers (Fig. 3b-c). Crucially, the sender and receiver weights were not uniformly distributed across the epithelial population; rather, they highlighted specific high-activity subpopulations, identifying the heterogeneity within this cell type (Fig. 3d-e). This subpopulation specificity was mirrored in the ligand-receptor usage: the top-weighted interactions—such as PTN-PTPRZ1, PTN-SDC1, COL4A5-SDC1, MDK-PTPRZ1, and MDK-SDC1—were not ubiquitously expressed but rather restricted to a fraction of epithelial cells (Supplementary Fig. 4a-c). Consequently, the highest communication scores were tightly localized to the cells most heavily weighted by the model (Fig. 3f). Normalized communication scores further confirmed that key signaling events were absent from the broader epithelial population (Supplementary Fig. 4d-f). Biologically, this implicates a distinct subset of cells in a pro-repair program, as PTN/MDK signaling via PTPRZ1/SDC receptors is known to reinforce tight junctions and support epithelial barrier integrity during host defense^38–40^.

**Figure 3.**
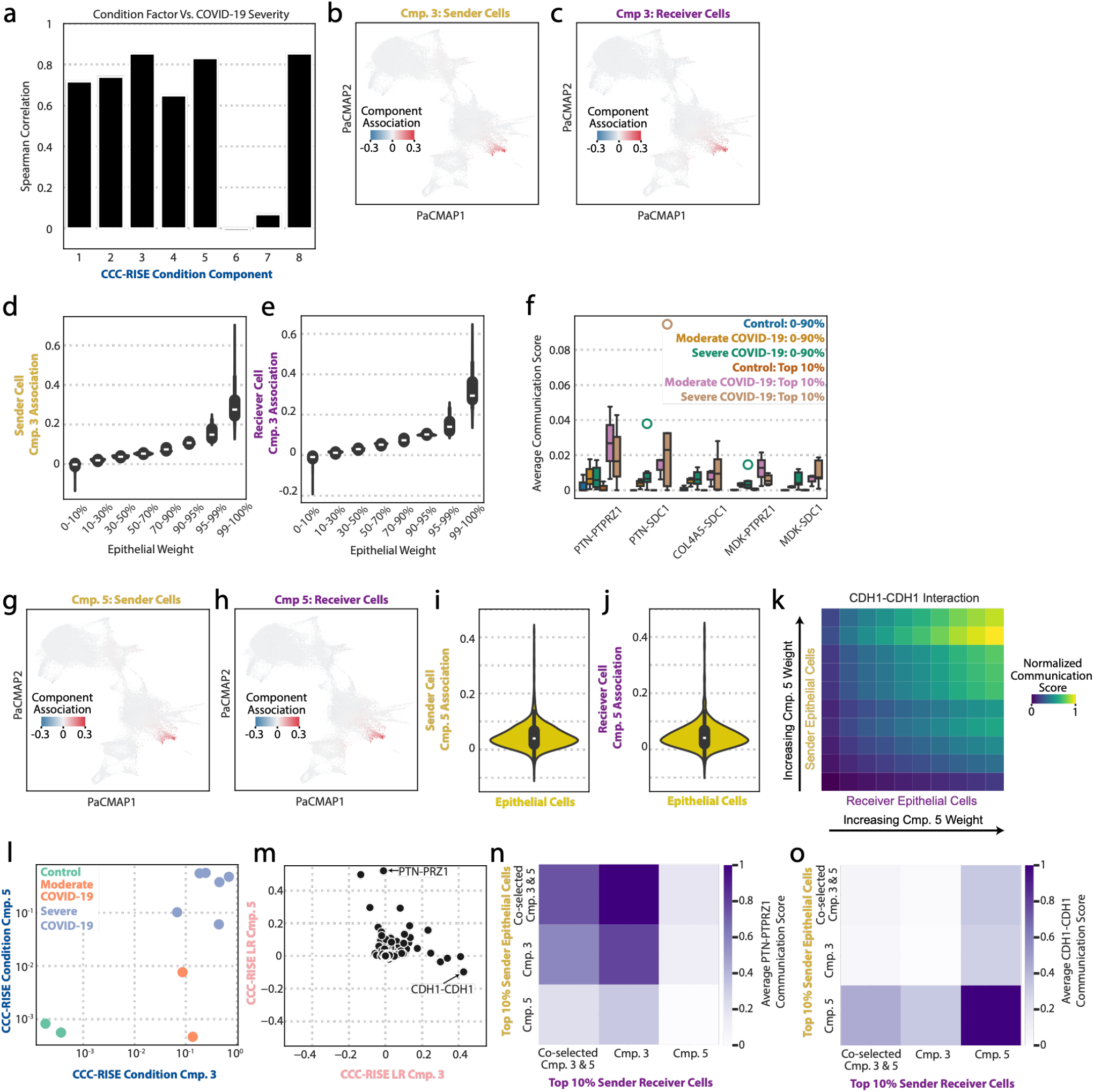
Identification of communication components correlated with COVID-19 severity. **(a)** Spearman correlation between CCC-RISE condition components and patient disease severity (ranked: healthy, moderate, severe). **(b, c, g, h)** PaCMAP embeddings of RISE projections visualizing single-cell heterogeneity. Cells are colored by their loadings for (b, g) sender and (c, h) receiver activity in (b-c) Component 3 and (g-h) Component 5. **(d, e, i, j)** Distribution of epithelial cell weights within (d, e) Component 3 and (i, j) Component 5, stratified by (d, i) sender and (e, j) receiver populations. **(f)** Average communication scores for the top five Ligand-Receptor (LR) pairs in Component 3, stratified by condition and cell weight. **(k, n, o)** Normalized communication scores for key interactions between top-weighted epithelial sender and receiver cells: (k, o) CDH1-CDH1 (Component 5) and (n) PTN-PTPRZ1 (Component 3 vs 5 comparison). **(l, m)** Comparison of Components 3 and 5, showing correlations between their (l) condition weights and (m) LR pair weights.

Like Component 3, Component 5 was strongly linked to disease severity and dominated by epithelial sender and receiver populations (Fig. 3g-h). However, it represented a distinct communication axis. The cellular weights again revealed intra-type heterogeneity, with specific subsets of epithelial cells driving the signal rather than the population as a whole (Fig. 3i-j). The molecular drivers here were different, characterized by adherence and junctional interactions such as CDH1-CDH1, OCLN-OCLN, and PRSS3-F2RL1 (Supplementary Fig. 4g). Communication scores for these pairs were notably variable; CDH1-CDH1 and OCLN-OCLN showed greater abundance compared to PRSS3-F2RL1, consistent with their higher weighting in the model (Fig. 3k, Supplementary Fig. 4h-i).

Despite both Component 3 and 5 being “epithelial-specific,” they described non-overlapping biological programs. They correlated with severity through distinct patient profiles (Fig. 3l) and utilized mutually exclusive sets of ligand-receptor pairs (Fig. 3m). This distinction extended to the cellular level (Fig. 3n-o). Cells heavily weighted in Component 3 showed strong PTN-PTPRZ1 communication but minimal involvement in Component 5’s programs (Fig. 3n). Conversely, the CDH1-CDH1 signal was robust specifically within the epithelial subset defined by Component 5 (Fig. 3o).

Finally, Component 8 also tracked with severity but displayed a broader connectivity pattern. While epithelial cells remained the primary senders, the signals—mediated largely by PTN-SDC3, PTN-SDC2, and PTN-SDC4 (Supplementary Fig. 4j)—were targeted toward a diverse array of receiver types, including B cells, mast cells, plasma cells, mDCs, and pDCs (Supplementary Fig. 4k–m).

### Distinct CD4+ T cell and NK cell signaling programs define ALAD

Having demonstrated CCC-RISE’s capacity to resolve subpopulation heterogeneity, we next evaluated its generalizability in a clinically distinct setting. We applied the framework to BAL samples from lung transplant recipients to distinguish patients with acute lung allograft dysfunction (ALAD) from stable controls (Fig. 4a). ALAD patients have a significant drop (10% decrease) in FEV_1_ (a measure of forced exhalation) and worsening oxygenation, leading to rapid lung function decline after transplantation and denoting concern for the onset of chronic lung allograft dysfunction (CLAD)^*41–43*^. The immune mechanisms driving this injury remain poorly understood^*44*^. Our cohort included a broad composition of immune and epithelial cell types; notably, there were no significant differences in the proportional abundance of these populations between ALAD and control patients, suggesting that the pathology is driven by altered communication rather than simple changes in cell numbers (Supplementary Fig. 5a). We optimized the CCC-RISE model using cross-validation and logistic regression, determining that a decomposition into 15 RISE eigen-states and 18 CPD components yielded the optimal predictive performance for ALAD status (Fig. 4b-c). The factors demonstrated high stability (Supplementary Fig. 5b-e) and effectively resolved distinct cell types within a common embedding space across all samples (Fig. 4f, Supplementary Fig. 5f).

**Figure 4.**
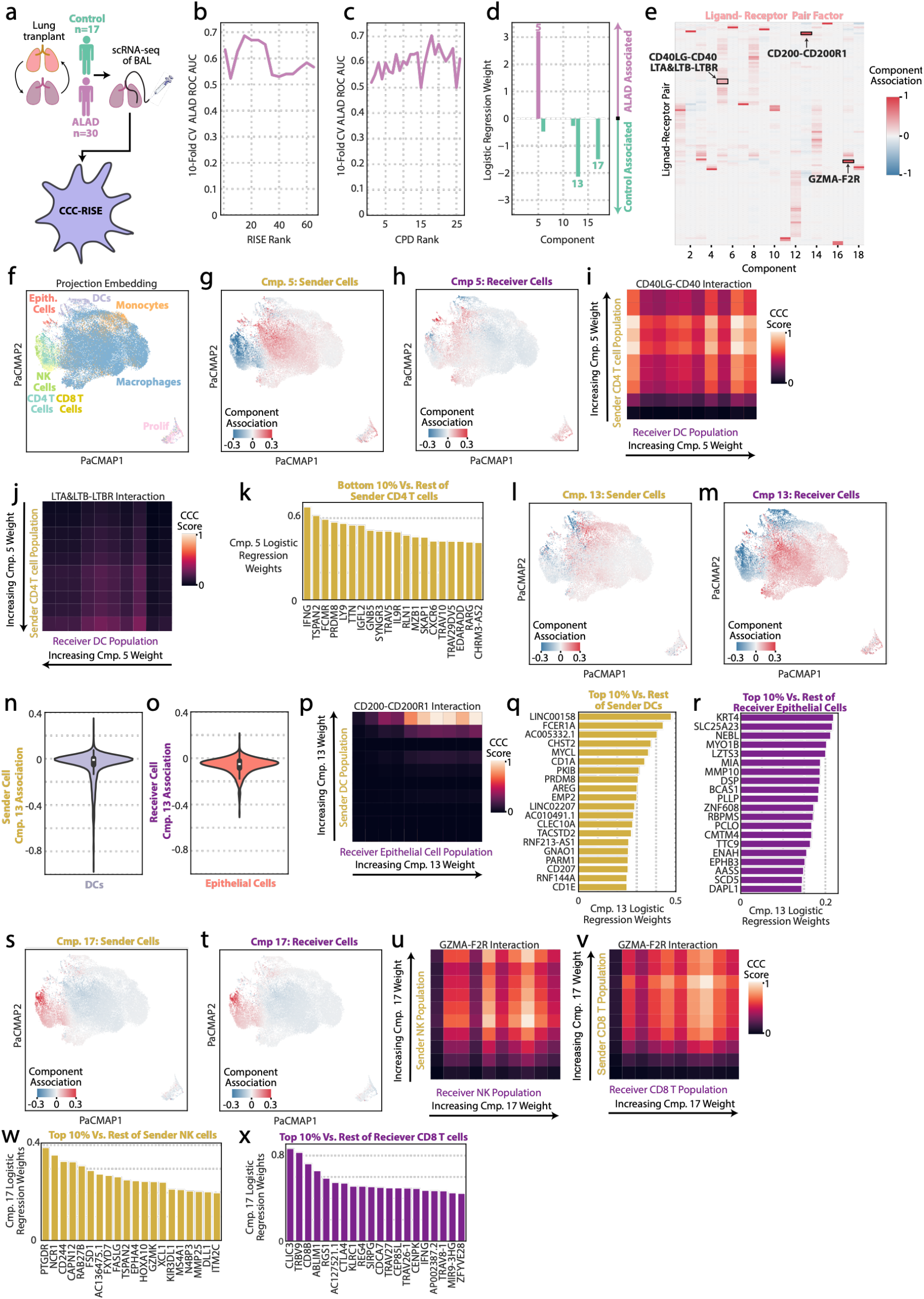
Identification of single-cell communication signatures associated with ALAD. **(a)** Workflow applied to BAL samples from lung transplant recipients to discriminate ALAD from control status. **(b-c)** ROC AUC performance of logistic regression classifiers using (b) RISE condition factors and (c) CPD condition factors across varying component ranks. **(d)** Feature weights for the optimal 18-component CPD logistic regression model; components 5, 13, and 17 are highlighted as top predictors. **(e)** Ligand-Receptor (LR) pair loadings for the top components. **(f)** PaCMAP embedding of RISE projections colored by cell type. **(g-h, l-m, s-t)** Visualization of key predictive components: PaCMAP embeddings show sender and receiver cell loadings for (g-h) Component 5 (ALAD-associated), (l-m) Component 13 (Control-associated), and (s-t) Component 17 (Control-associated). **(i-j, p, u-v)** Normalized communication scores for top interactions: (i-j) Component 5-driven CD40LG-CD40 and LTA-LTBR signaling between CD4+ T cells and DCs; (p) Component 13-driven CD200-CD200R1 signaling between DCs and epithelial cells; and (u-v) Component 17-driven GZMA-F2R signaling among NK and CD8+ T cell populations. **(k, q-r, w-x)** Differentially weighted genes based on logistic regression distinguishing the (k) bottom 10% from the remaining sender CD4+ T cells for Cmp. 5, the bottom 10% (q) sender DCs for Cmp. 13 and (r) receiver epithelial cells for Cmp. 13, and the top 10% (w) sender NK cells and (x) receiver CD8 T cells for Cmp. 17. **(n-o)** Distribution of Component 13 loadings within (n) DCs and (o) epithelial cells.

The logistic regression analysis identified three primary components that strongly discriminated between patient groups: Component 5 was associated with ALAD status, while Components 13 and 17 were associated with control samples (Fig. 4d). These associations remained robust even when analyzing pairwise combinations of components, indicating they represent stable biological signals (Supplementary Fig. 5g). The model successfully captured distinct communication programs for each component, defined by specific weights across LR pairs and cell populations (Fig. 4e, Supplementary Figs. 5h, 6, and 7).

Investigating the ALAD-associated signal, Component 5 was characterized by a specific interaction between CD4+ T cells and dendritic cells (DCs) (Fig. 4g-h). This program was dominated by CD40LG-CD40 signaling, which exhibited a much stronger interaction profile and weighting compared to other pairs like LTA-LTBR (Fig. 4e, 4i-j). Crucially, the communication scores for this interaction revealed considerable heterogeneity within the CD4+ T cell and DC populations, confirming that the signal is driven by specific active subsets rather than the total population. Examining the transcriptional signature of CD4+ T cells with greater communication revealed features suggestive of regulatory T cells (FOXP3, TNFRSF25) with stem-like features (SELL, LEF1) participating in lymphoid-like structures (LTA; Fig. 4k).

In contrast, Components 13 and 17 defined distinct signaling networks associated with stable control patients. Component 13 was driven primarily by interactions between DCs and epithelial cells (Fig. 4l-o). The top-weighted LR pairs for this component included CD200-CD200R1 and IL27-related interactions, which were distinct from those found in other components (Fig. 4e, Supplementary Fig. 5i). Communication scores highlighted a prominent role for DCs and epithelial cells in this potentially protective program, particularly via the CD200-CD200R1 axis (Fig. 4p). DCs with greater sender activity had features of Langerhans cells (CD207, CD1A, CD1E, CD301), while epithelial cells with greater receiver activity had associations with a unique differentiation program (e.g. KRT4, DSP, MMP10) (Fig. 4q-r). Finally, Component 17 revealed a separate control-associated program involving NK cells and CD8+ T cells (Fig. 4s-t). This component was heavily weighted toward the GZMA-F2R interaction (Fig. 4e), with single-cell resolution analysis confirming that this signaling axis is actively utilized by both NK and CD8+ T cell populations in stable grafts (Fig. 4u-v). High communication NK cells were enriched for other transcriptional features associated with activation and cytotoxicity, such as FASLG and 2B4, while the high receiving CD8+ T cells showed other markers of activation, such as CTLA4 and CLIC3 (Fig. 4w-x).

### CCC-RISE identifies communication signatures of long-term clinical stability after ALAD

Having defined the cellular communication networks associated with ALAD, we sought to contextualize their broader clinical significance by comparing the identified CCC-RISE components against patient demographic and physiological variables. We found no significant correlations between the components and variables such as sex, age, or FEV_1_ measured at 6 months and 1-year post-transplant (Supplementary Fig. 8a-b). Beyond the initial ALAD diagnosis, we investigated whether these components could predict progression to CLAD or mortality, outcomes marked by continued functional decline^*41*^. Our analysis revealed that the sample associations for these components did not clearly associate with either CLAD or deceased status at 6 months and 1-year post-ALAD diagnosis as separate outcomes (Supplementary Fig. 8c-h).

Given these findings, we expanded our analysis to determine if other signatures from CCC-RISE could stratify patients based on their clinical trajectories, specifically distinguishing those who remained stable from those who experienced clinical decline leading to CLAD or death in aggregate. Using the original factorization, we applied logistic regression to identify components predictive of these outcomes. Components 7 and 17 emerged as significant predictors for stratifying patient stability (Fig. 5a). Notably, the components differentiating stable status were consistently identified as significant when assessing pairwise interactions, even if they were potentially filtered out by L1 regularization in isolation (Supplementary Fig. 8i).

**Figure 5.**
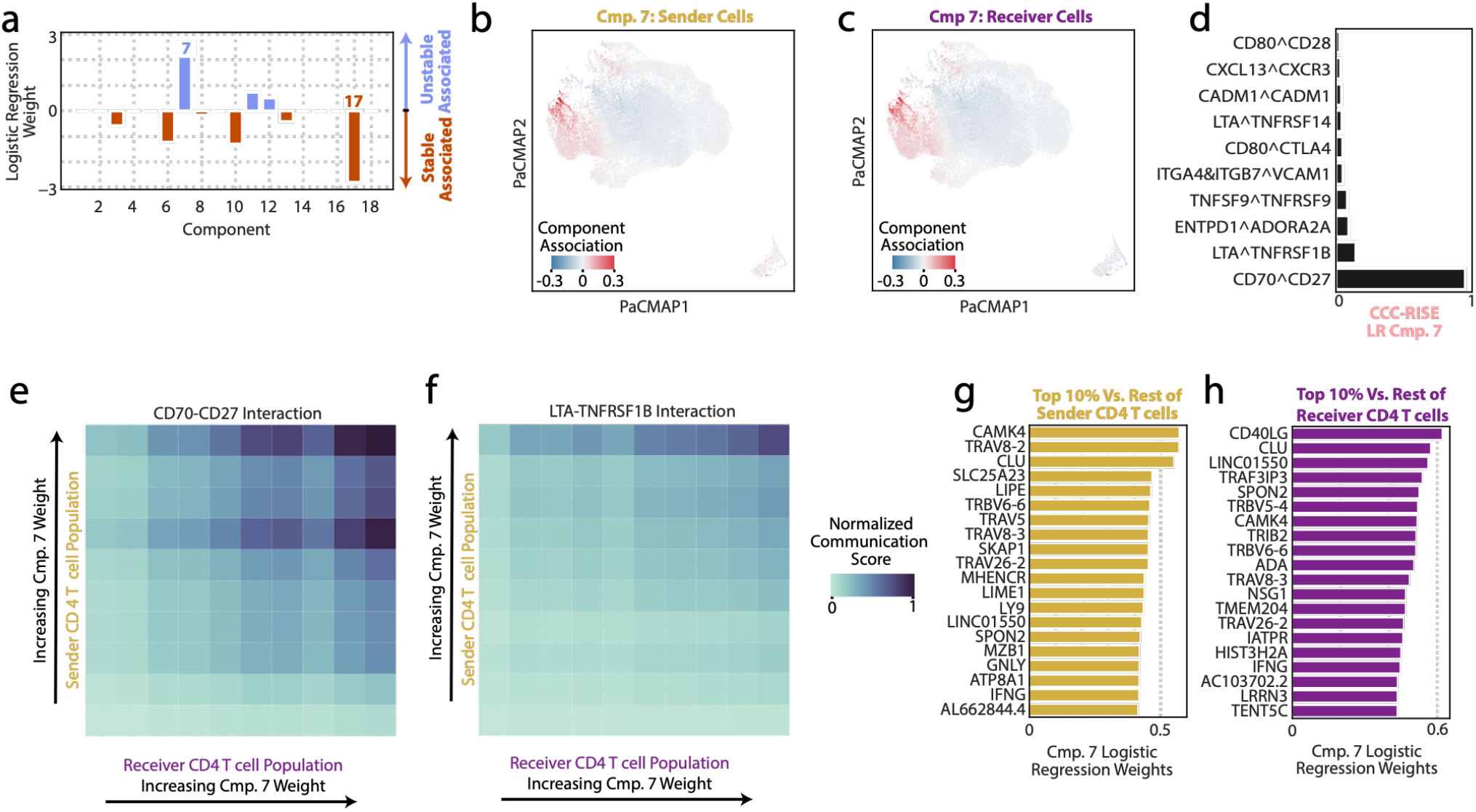
Communication programs distinguish stable outcomes from chronic decline in lung transplants. **(a)** Logistic regression weights for an 18-component CPD model, identifying the most predictive components for patient stability. **(b-c)** PaCMAP embeddings of RISE projections, with (b) sender and (c) receiver cells colored by their loadings in Component 7. **(d)** The top 10 most positively weighted ligand-receptor (LR) pairs identified by CCC-RISE for Component 7. **(e-f)** Normalized communication scores between Component 7 sender CD4+ T cells and receiver CD4+ T cells for the (e) CD70-CD27 and (f) LTA-TNFRSF1B signaling axes. **(g-h)** Differentially weighted genes based on logistic regression distinguishing the top 10% (g) sender NK cells and (h) receiver CD4 T cells for Cmp. 7.

The analysis reaffirmed the importance of the GZMA-F2R and CCL5-F2R interactions within NK and CD8+ T cell populations as associated with better outcomes, this time stable lung function after ALAD. These interactions were previously identified as predictors for control patients without ALAD, suggesting that the GZMA-F2R axis represents a consistent communication pathway in successful allografts, likely reflecting appropriate immune surveillance that avoids excessive inflammation. Conversely, Component 7 was the most heavily weighted predictor of clinical decline, highlighting a distinct signal driven by CD4+ T cells (Fig. 5b-c). The strongest ligand-receptor interactions included CD70-CD27 and LTA-TNFRSF1B (Fig. 5d). Detailed analysis of CD4+ T cell communication patterns revealed that the CD70-CD27 interaction profile was substantially more robust than that of LTA-TNFRSF1B or other pairs (Fig. 5e-f). Analysis of the distinguishing features of senders and receivers identified overlapping markers of activation (CAMK4, CLU; Fig. 5g-h). This implicates the CD70-CD27 axis as a potential driver of pathological signaling associated with clinical deterioration. CD70+ CD4+ T cells were similarly identified in an independent cohort of small airway brushings from recipients with stable allograft function or ALAD^*44*^. Despite the limited sample size, the fraction of CD70+ CD4+ T cells were elevated in ALAD compared to stable controls (Supplementary Fig. S9).

## DISCUSSION

We have demonstrated that CCC-RISE provides a comprehensive framework for investigating cell-cell communication (CCC) across multiple conditions while maintaining single-cell resolution. Current computational tools for CCC largely rely on pseudobulking, a process that aggregates single-cell data into pre-defined clusters and obscures the functional diversity of individual cells. This approach depends heavily on the accuracy of initial annotations and often fails to capture the continuous and multifaceted nature of cellular identities. To address these limitations, CCC-RISE leverages RISE to project cells into data-driven “eigen-states” based on shared expression patterns rather than rigid cell types. By applying tensor decomposition to these eigen-states, CCC-RISE identifies integrative patterns of variation, allowing researchers to trace communication programs back to specific sender and receiver subpopulations, even those that cross conventional cell-type boundaries.

The application of CCC-RISE to a COVID-19 cohort illustrated the critical need for this resolution to deconvolve condition-specific communication events. While pseudobulk analysis broadly implicated mDCs and B cells in CCL19-CCR7-mediated interactions, CCC-RISE successfully pinpointed the specific high-activity subpopulations driving this recruitment signal, revealing heterogeneity that aggregate methods miss. Furthermore, the model disentangled complex epithelial responses by identifying a distinct pro-repair program involving PTN/MDK signaling through PTPRZ1/SDC receptors^45–47^. This signaling axis was associated with only a fraction of the epithelial cell population, aligning with established host defense mechanisms where specific proteins like occludin and E-cadherin are upregulated to reinforce tight junctions and prevent viral transmission^48,49^. The ability to distinguish these non-overlapping ligand-receptor programs within a single “cell type” highlights the method’s capacity to separate disease mechanisms that are mathematically invisible to aggregate approaches.

We further validated the utility of CCC-RISE in a cohort of lung transplant patients by identifying specific communication programs associated with the onset of ALAD. Crucially, the framework demonstrated unique capacity to resolve communication at a subpopulation level, uncovering functional heterogeneity within standard cell type annotations. In ALAD patients, CCC-RISE pinpointed pathogenic CD40LG-CD40 signaling between dendritic cells and a specific subset of CD4+ T cells enriched for stem-like, homing, and regulatory markers, including FOXP3, SELL, and LEF1. This distinct transcriptional profile suggests these cells actively organize the immune response within lymphoid-like structures, a critical biological nuance that aggregate methods would miss. Indeed, markers of lymphoid structure formation and regulation have previously been identified as associated with acute cellular rejection^50^. Conversely, CCC-RISE also identified counterbalancing, pro-tolerance communication programs. First, DC interactions for CD200-CD200R1 suggest a communication program where DCs engage in an anti-inflammatory response and promotes immune tolerance, as suggested by previous studies^51–53^. Secondly, GZMA-F2R was the primary interaction for NK cells and terminally exhausted CD8 T cells previously identified as predictive of successful outcomes in control patients and conducive to transplant tolerance, potentially reflecting effective immune surveillance without detrimental inflammation. By capturing the simultaneous expression of cytotoxic machinery and inhibitory checkpoints like CTLA4 in these receivers, the framework successfully identified a state of strictly regulated immune surveillance that prevents detrimental inflammation. Ultimately, these findings highlight that complex clinical outcomes like ALAD are associated with the signaling of highly specialized, transient cell subsets, underscoring the necessity of integrative analytical tools that can bypass rigid cell annotations.

Building on the characterization of acute dysfunction, we used CCC-RISE to identify communication networks that predict long-term clinical trajectories. The model identified Component 7 as the strongest predictor of clinical instability, driven almost entirely by an aberrant CD70-CD27 signaling axis within the CD4+ T cell compartment. While CD70 is typically repressed shortly after normal T cell activation, sustained expression marks alloreactive T cells^54^. The single-cell resolution of CCC-RISE isolated this specific subpopulation of CD4+ T cells characterized by high CD70 expression. By engaging CD27 on neighboring cytotoxic CD4+ receiver cells—which display distinct, high-metabolic markers such as ADA—these CD70+ senders create a continuous, self-amplifying feedback loop. This aligns with the known role of CD70-CD27 in promoting antigen-driven T-cell proliferation and lowering the activation threshold of the surrounding immune environment^54–57^. Mathematically separating this dysregulated, highly specialized subset from the broader CD4+ population again demonstrated the importance of cellular resolution beyond the cell type annotations.

A central advantage of this framework is its ability to bypass the limitations of cell annotations. By analyzing eigen-states, CCC-RISE captures continuous and overlapping cellular identities, which is particularly critical for rare cell populations or for identifying subpopulations that participate in multiple, distinct signaling programs. However, despite these advantages, the method has limitations inherent to data reduction and transcriptomic inference. Users must define the number of eigen-states and components, a flexible but challenging process that can be aided by metrics such as the Factor Match Score^21,22^. Future work could incorporate subsampling approaches to assess factor reproducibility and enable empirical power calculations to estimate the sample size required to detect meaningful variation^58–61^. Additionally, like all transcriptomic inference tools, CCC-RISE relies on the assumption that gene expression is a sufficiently accurate surrogate of protein abundance and functional interaction. Because post-transcriptional regulation and binding kinetics can decouple mRNA levels from functional protein interactions, the communication patterns identified should be interpreted as hypotheses requiring experimental validation^8,62^. Emerging technologies like single-cell proteomics will be crucial for CCC inferring methods to improve the accuracy of predictions^63,64^.

In summary, CCC-RISE provides an unbiased framework for investigating CCC across multiple conditions at single-cell resolution. By integrating the cell-alignment capabilities of RISE with the pattern-finding benefits of tensor decomposition, our method reveals context-specific communication programs that are directly interpretable at the level of individual cells. As single-cell multi-condition studies become increasingly common, CCC-RISE serves as a powerful tool to unravel the complex cellular dialogues that underpin diseases. Understanding CCC will rely on high-level analysis tools like CCC-RISE to map the entire landscape of potential interactions in an unbiased way, followed by targeted experiments to confirm which of these predicted pathways are functionally important.

## Acknowledgements

This work was supported by NIAID U19-AI172713 to A.S.M. and a Cota Robles Fellowship to A.R. We thank Erick Armingol and Hratch M. Baghdassarian for their invaluable advice for the development of this method and its application to cell-cell communication.

## Author Contributions

A.R. and A.S.M. conceived of the study. All authors performed the computational analysis and helped analyze the data. All authors wrote and revised the paper.

## Supplemental Information

Supplemental Information is attached including Supplementary Figures 1–8.

## Resource Availability

### Lead Contact

Further information and requests for resources should be directed to and will be fulfilled by the Lead Contact, Aaron Meyer.

### Materials Availability

This study did not generate new unique reagents.

### Data and Code Availability

- All analysis was implemented in Python v3.12 and can be found at https://github.com/meyer-lab/cellcommunication-Pf2, release 1.0.
- The implementation of CCC-RISE is available at https://github.com/meyer-lab/cellcommunication-Pf2. The single-cell gene expression was obtained from Calabrese et al^41^.
- Any additional information required to reproduce this work is available from the Lead Contact.

## MATERIALS AND METHODS

### RISE, CPD, and CCC-RISE

#### Canonical Polyadic Decomposition

Canonical Polyadic Decomposition (CPD) is a tensor decomposition method that generalizes PCA to higher-order data structures. While PCA operates on matrices (second-order tensors), CPD extends this logic to third-order (and higher-order) tensors. Given a third-order tensor, ***X***, of shape *I* × *J* × *K*, CPD seeks to approximate ***X*** as a sum of component rank-one tensors. This set of component rank-one tensors are collectively represented in factor matrices: *A* of shape *I* × *R, B* of shape *J* × *R*, and *C* of shape *K* × *R*, where *R* is the number of components into which ***X*** is decomposed. Specifically, CPD seeks

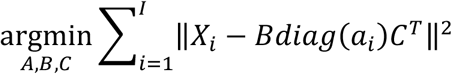

where *X*_***i***_ is the *i*^th^ slice of ***X*** along the first mode, and *diag*(*a*_*i*_) is the diagonal matrix whose nonzero entries are equal to the *i*^th^ row of *A*. While many algorithms exist for deriving CPD, we applied alternating least squares (ALS), wherein each mode is iteratively solved using least squares. A tolerance of 1×10^-11^ and maximum of 10,000 iterations was defined. As an iterative procedure, ALS must be initialized with a starting estimate of the factorization. The factor matrices were initialized using the SVD of the matrix-flattened data. Each of *X*_*i*_ was concatenated to form a matrix. Randomized SVD was performed on this matrix and the first *R* right singular vectors were used to initialize *B. A* was filled with ones and *C* was initialized to the identity matrix.

#### Reduction and Insight in Single-cell Exploration

Reduction and Insight in Single-cell Exploration (RISE) is a computational framework based on the PARAFAC2 tensor decomposition method, optimized for analyzing multi-condition scRNA-seq data. A major challenge in scRNA-seq is that the data lacks alignment when reconstructed into a 3D tensor, because the same cell is not measured across conditions. RISE overcomes this by using the PARAFAC2 model, which solves for the alignment of the data along one dimension—the cell axis. In fitting the RISE model, we seek:

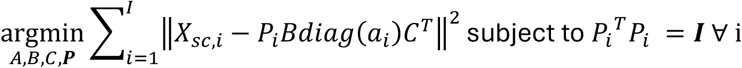

where 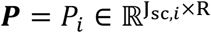, i = 1, 2, …, I are the projection matrices and *R* is the rank of the RISE decomposition; *diag*(*a*_*i*_) is the diagonal matrix whose nonzero entries are equal to the *i*^th^ row of *A* ∈ ℝ^*I*×*R*^, the condition factor matrix; *B* ∈ ℝ^*R*×*R*^ is the cell eigen-state factor matrix; *C* ∈ ℝ^*K*×*R*^ is the gene factor matrix; and ***I*** is the identity matrix^65^. We fit RISE to obtain ***P*** and the factor matrices, *A, B*, and *C*, which are alternately fit. First, the factor matrices are updated as

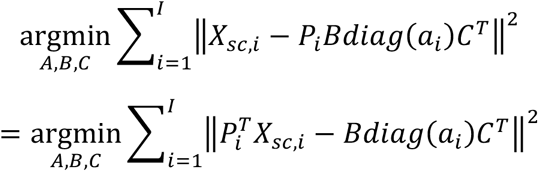

which is equivalent to running PARAFAC on the single, aligned, third-order tensor whose i^th^ slice along the first mode is equal to 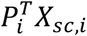. Then, the projection matrices, ***P***, are updated by solving the Orthogonal Procrustes problem. Given matrices X, Y ∈ ℝ^n×m^, the Orthogonal Procrustes problem seeks an orthogonal matrix *Q* ∈ ℝ^*n*×*m*^ that minimizes 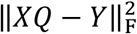. The solution is given by Q = UV^T^, where UΣV^T^is the singular value decomposition of Y^T^X. For a low-rank decomposition, we update each **P**_***i***_ ∈ ***P*** as the product of the first *R* left and right singular vectors from the SVD of 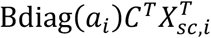. This process of updating the factor matrices using PARAFAC and the projections via Orthogonal Procrustes is repeated until a specific number of iterations elapses or the difference in R2X between iterations reaches a threshold. This cycle of aligning the tensor and then factoring it is repeated until either 100 iterations elapse or the difference in R^2^X between iterations falls below 1×10^−6^. RISE enhances the core PARAFAC2 algorithm with several key optimizations including a parameter initialization scheme via SVD, the implementation of Nesterov acceleration to improve convergence, and post-factorization alignment procedures to ensure consistent, interpretable results.^65,66^

#### CCC-RISE

CCC-RISE is a computational pipeline for the analysis of CCC across multiple conditions from scRNA-seq data. The method proceeds in three key steps. The scRNA-seq data tensor is first decomposed using RISE. From the eigen-state tensor that RISE produces based on the factors, CCC scores are calculated for all relevant LR pairs in each experimental condition, with each pair of eigen-states as sender and receiver.

To model CCC, we represent the expression of a ligand and its receptor as the outer product of the gene expression for a LR pair. This operation captures the combinatorial potential for signaling, where each element in the resulting matrix represents the product of a sender eigen-states’ ligand expression and a receiver eigen-states receptor expression. In doing so, we assume the LR interactions are proportional to the mRNA abundance of the ligand and receptor. When extended across all ligand-receptor pairs and conditions, these form an aligned 4D communication tensor with dimensions of sender eigen-state, receiver eigen-state, LR pair, and condition. CPD is then applied to the 4D communication tensor, resulting in factor matrices that explain the component associations with each of the four dimensions.

### Data Normalization, Processing, and Visualization

#### BAL COVID-19

The single-cell gene expression was obtained from Liao et al^33^. Genes with less than 0.001 mean counts across all cells were removed. The size of the dataset included 63,103 cells and 17,714 genes. Each cell was then normalized by read depth using CPM normalization. Next, each gene was scaled according to its sum. Finally, the data was log transformed via the Delta method,

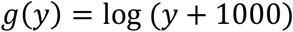

where *g*(*y*) is the transformed gene and *y* is the pre-transformed gene. Finally, each gene was mean-centered. The data was then fit using the RISE model with 35 components as described above. PaCMAP was applied to the concatenated matrix of the projections across all conditions to visualize cell-to-cell variation as an embedding. PaCMAP’s default parameters of n_neighbors=10, MN=0.5, and FP_ratio=2.0 were used^67^. The communication tensor was calculated based on a human list of 2,005 ligand-receptor pairs, including multimeric protein complexes, was obtained from CellChat^5^. We filtered this list by considering the genes expressed in the dataset, resulting in 895 LR pairs. The data was then fit using the CPD model with 8 components as described above. Cell annotations were reused from the initial data collection and analysis^33^.

#### BAL ALAD

The single-cell gene expression was obtained from Calabrese et al^41^. The same normalization pipeline was used as described above. The final dataset consisted of 51,273 cells across all conditions and 15,899 genes. The data was first fit using the RISE model with 15 components. For visualization, the same PaCMAP parameters were applied. After filtering the list and considering the genes expressed in the dataset, a total of 445 LR pairs were identified. The data was then fit using the CPD model with 18 components. Cell annotations were performed during the initial data collection and subsequent analysis^41^.

### Reconstruction Error

Reconstruction error (R2X), or the fraction of variance explained by the model, was used to evaluate the effectiveness of the data reduction. For RISE, the Frobenius norm of the difference between the scRNA-seq data (X_data_) and the reconstruction from the RISE factors and projections (X_RISE_) is calculated. This difference is normalized by the Frobenius norm of the data and subtracted from 1.

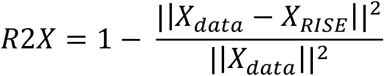

For CPD, the Frobenius norm of the difference between the 4D communication tensor after CCC scoring (X_CCC 4E Tensor_) and the reconstruction from the CPD factors (X_CPD_) is calculated. This difference is normalized by the Frobenius norm of the data and subtracted from 1:

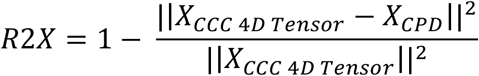

R2X ranges from 0 to 1, with a score of 1 indicating a perfect reconstruction.

### Factor Match Score

The factor match score (FMS) was used to evaluate the stability of the RISE and CPD factorization^68^. First, the condition (*A*) and gene (*B*) factors were derived for RISE. To generate a comparable dataset, cells were resampled with replacement to produce a bootstrapped dataset with an equal number to the original dataset with a different composition. RISE factorization was performed on this bootstrapped dataset with the same number of components, and the bootstrapped condition (*Â*) and gene 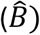 factors were collected. Only the condition (*A, Â*) and gene 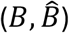 factor matrices were compared between factorizations; cell eigen-state factors were not compared as the cells used in the bootstrapped factorization are different from those in the original factorization.

Component orders in RISE are not fixed: two factorizations can have equivalent components in different orders. As such, linear sum assignment was applied to align components between factor matrices. Following component alignment, FMS is derived as follows:

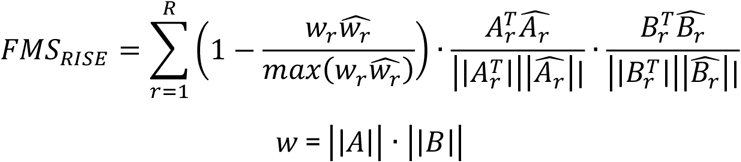

This process was repeated a total of five times to produce five sets of bootstrapped factors. FMS was derived between the base factors and each set of bootstrapped factors to produce five FMS scores. FMS scores range between 0 and 1 with a score of 1 indicating that two sets of factors are identical.

After the number of components were chosen for RISE and the communication tensor was calculated, LR pairs were resampled with replacement to produce a bootstrapped dataset with an equal number to the original dataset with a different composition. CPD factorization was performed on this bootstrapped tensor with the same number of components, and the bootstrapped condition (*Ĉ*), sender eigen-state 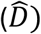, and receiver eigen-state (*Ê*) factors were collected. Only the condition (*C, Ĉ*), sender eigen-state 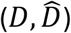 and receiver eigen-state (*E, E*) factor matrices were compared between factorizations.

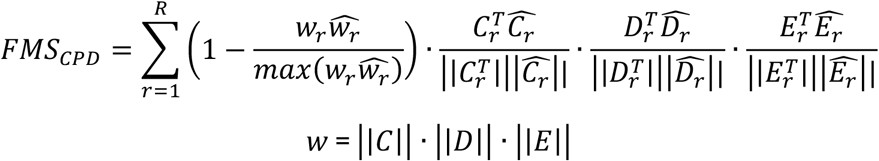

This process was repeated a total of five times to produce five sets of bootstrapped factors. FMS was derived between the base factors and each set of bootstrapped factors to produce five FMS scores.

### Logistic Regression

Regularized logistic regression was implemented using scikit-learn, with an L1 penalty, the Stochastic Average Gradient (SAGA) solver, a tolerance of 1×10^-9^, and maximum iteration number of 10,000^69^. The regularization strength was determined through cross-validation using the LogisticRegressionCV and the default parameters regarding attempted regularization strengths. We applied repeated, stratified, 10-fold cross-validation, with 5 repeats throughout the analysis.

### Cell Type Composition and Pseudobulking

Cell type composition was analyzed by calculating the percentage of cells belonging to each cell type within an experimental condition. Pseudobulk profiles were created by averaging the normalized expression for each cell type in each condition.

### Quantification and Statistical Analysis

Figure captions detail pertinent statistical analyses or metrics, the number of conditions, and confidence interval values.

