## Supplementary Materials for "Tracing cell communication programs across conditions at single cell resolution with CCC-RISE"

### SUPPLEMENT

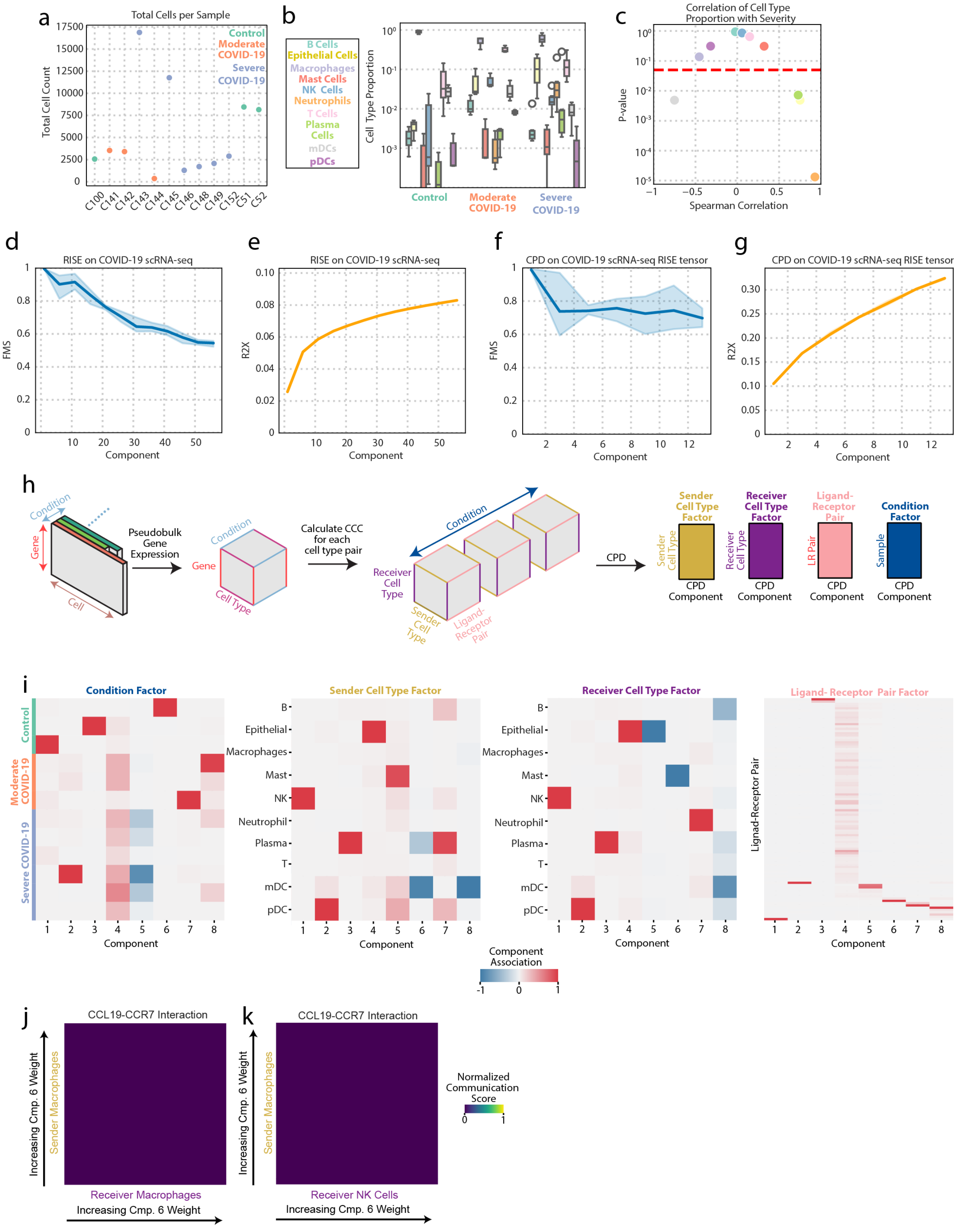

a
Total Cells per Sample
Total Cell Count
Control
Moderate COVID-19
Severe COVID-19
C100 C141 C142 C143 C144 C145 C146 C148 C149 C152 C51 C52
b
B Cells
Epithelial Cells
Macrophages
Mast Cells
NK Cells
Neutrophils
T Cells
Plasma Cells
mDCs
pDCs
Cell Type Proportion
Control
Moderate COVID-19
Severe COVID-19
c
Correlation of Cell Type Proportion with Severity
P-value
Spearman Correlation
d
RISE on COVID-19 scRNA-seq
FMS
Component
e
RISE on COVID-19 scRNA-seq
R2X
Component
f
CPD on COVID-19 scRNA-seq RISE tensor
FMS
Component
g
CPD on COVID-19 scRNA-seq RISE tensor
R2X
Component
h
Condition
Gene
Cell
Pseudobulk Gene Expression
Cell Type
Calculate CCC for each cell type pair
Receiver Cell Type
Sender Cell Type
Ligand-Receptor Pair
CPD
Sender Cell Type Factor
Receiver Cell Type Factor
Ligand-Receptor Pair
Condition Factor
LR Pair
Sample
CPD Component
i
Condition Factor
Sender Cell Type Factor
Receiver Cell Type Factor
Ligand- Receptor Pair Factor
B
Epithelial
Macrophages
Mast
NK
Neutrophil
Plasma
T
mDC
pDC
Ligand-Receptor Pair
Component
Component Association
j
CCL19-CCR7 Interaction
Increasing Cmp. 6 Weight
Sender Macrophages
Receiver Macrophages
k
CCL19-CCR7 Interaction
Receiver NK Cells
Normalized Communication Score

**Figure S1. CCC-RISE model validation and COVID-19 dataset characterization. (a)** Cell type proportion across the patient categories (B cells, epithelial cells, macrophages, natural killer cells, neutrophils, T cells, plasma cells, myeloid-derived suppressor cell, and plasmacytoid dendritic cells. **(b)** Total cell count per sample with points labeled by their COVID-19 status. **(c)** Spearman between cell type proportion and the severity of patient samples (1: healthy, 2: moderate, 3: severe). **(h)** CCC analysis framework using pseudobulk gene expression based on cell types and the tensor factorization CPD. **(d)** Factor match score (FMS) when comparing the RISE gene and condition factors of the full scRNA-seq COVID-19 dataset to a bootstrapped dataset at various ranks. **(e, g)** Fraction of the variance explained by (b) RISE and (d) CPD for each number of components**. (f)** FMS when comparing the CPD condition and sender/receiver eigen-state factors of the 4D communication eigen-state tensor (35-component RISE model) to a bootstrapped tensor at various ranks. **(i)** The condition, sender cell type, receiver cell type, and LR pair factors for each component in the reduced pseudobulk communication tensor. **(j-k)** Normalized communication score between the most weighted sender macrophages and the most weighted (j) receiver macrophages and (k) NK cells for CCL19-CCR7 in for component 6.

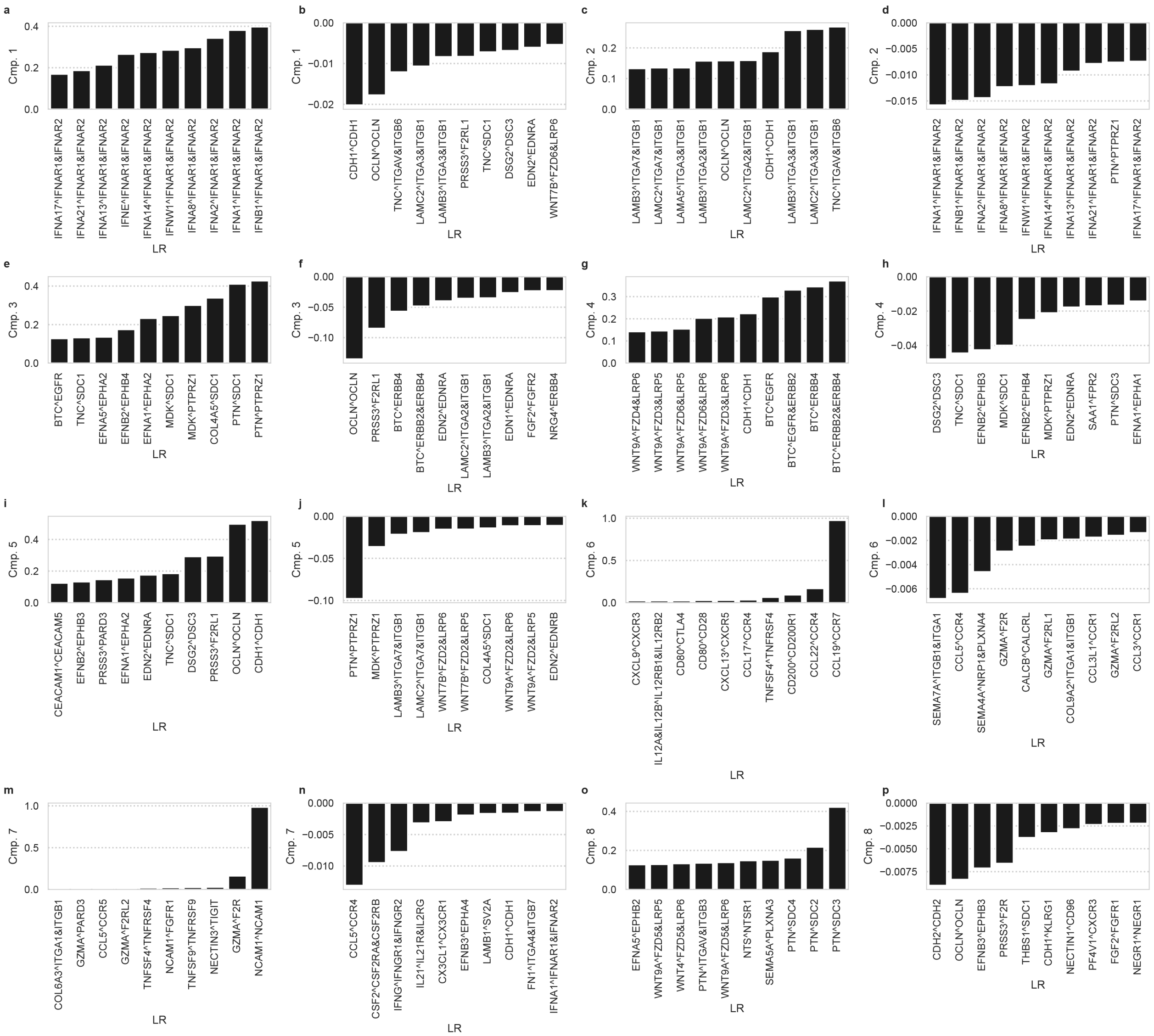

**Figure S2. CCC-RISE components associate each pattern to specific LR pairs for the scRNA-seq COVID-19 dataset. (a, c, e, g, i, k, m, o)** The top 10 most positively weighted LR pairs identified by CCC-RISE by a 35-component RISE and 8-component CPD model. **(b, d, f, h, j, l, n, p)** The top 10 most negatively weighted LR pairs identified by CCC-RISE by a 35-component RISE and 8-component CPD model.

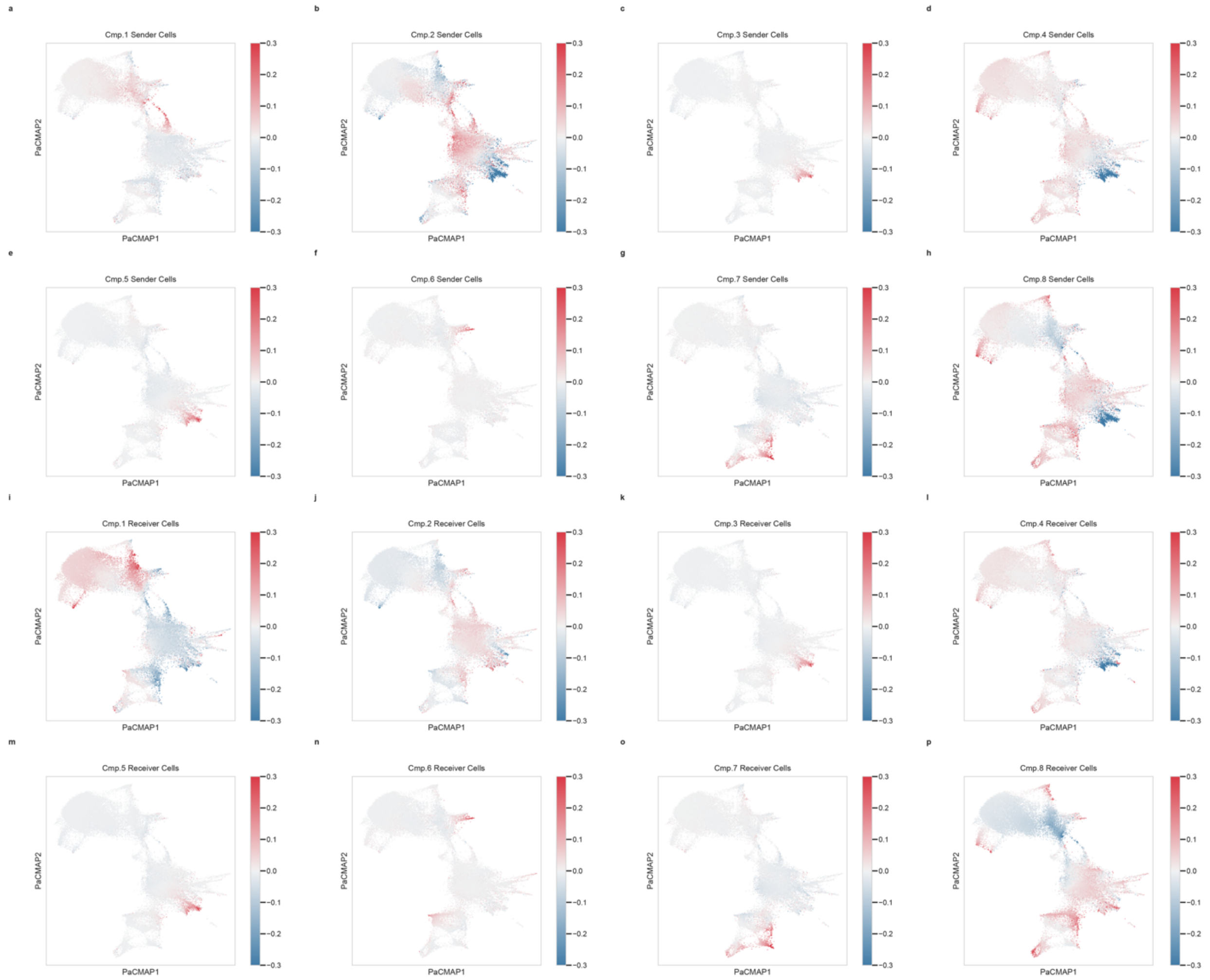


**Figure S3. CCC-RISE components associate each pattern to specific sender and receiver cells for the scRNA-seq COVID-19 dataset for a 35-component RISE and 8-component CPD model. (a-p)** PaCMAP of the RISE projections, with (a-h) sender cells and receiver cells (i-p) colored by the weightings for the indicated component.

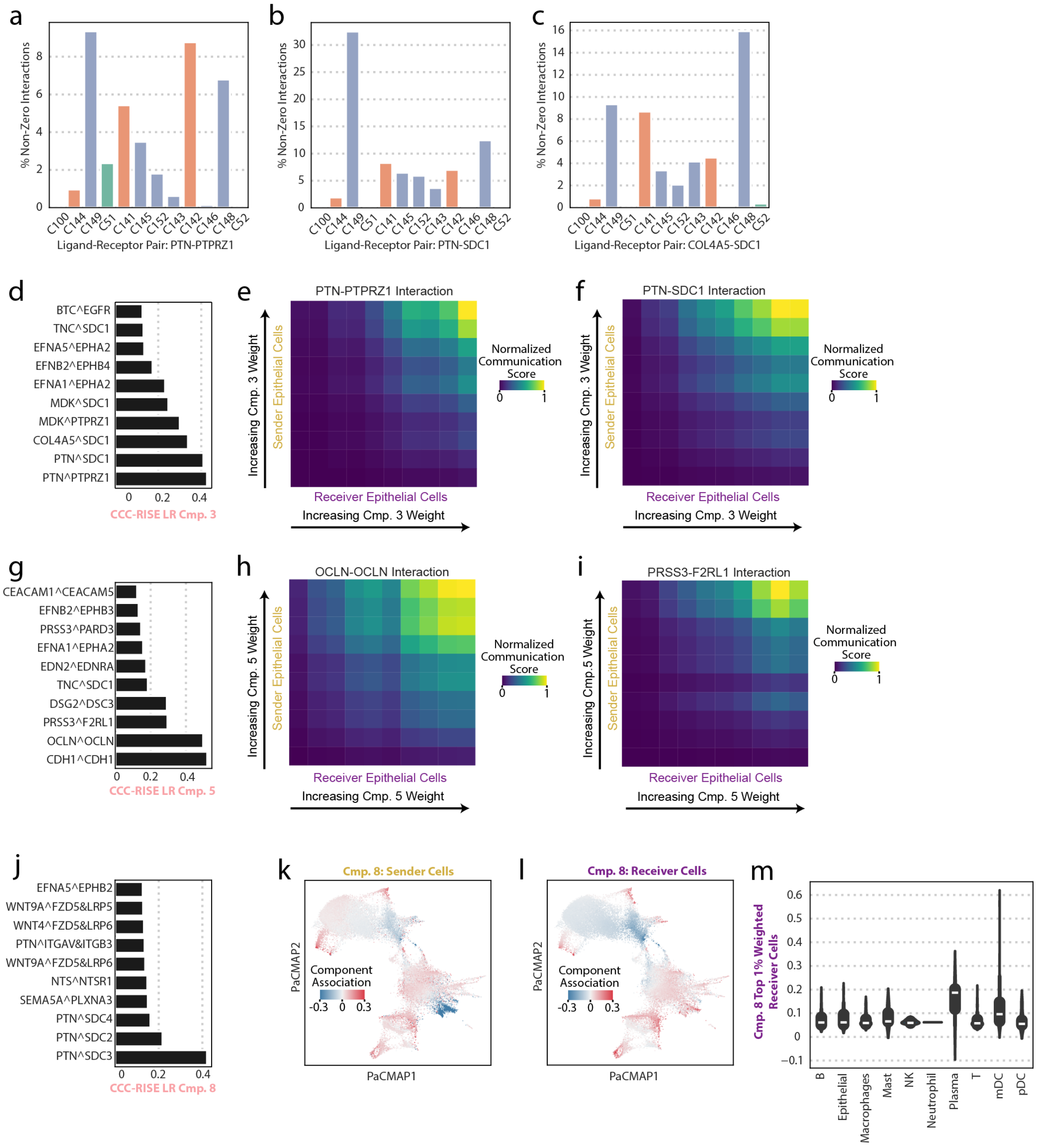


**Figure S4. CCC-RISE captures biologically relevant cell-cell communication associated with mortality. (a-c)** Percentage of non-zero interactions for all epithelial cells for each sample in (a) PTN-PTPRZ1, (b) PTN-SDC1, and (c) COL4A5-SDC1. **(d, g, j)** The top 10 most positively weighted LR pairs identified by CCC-RISE for (d) component 3, (g) component 5, and (k) component 8. **(e-f, h-i)** Normalized communication score between the most weighted epithelial cells for (e) component 3 and PTN-PTPRZ1, (f) component 3 and PTN-SDC1, (g) component 5 and OCLN-OCLN, and (e) component 5 and PRSS3-F2RL1. **(k-l)** PaCMAP of the RISE projections, with (k) component 8 sender cells and (l) component 8 receiver cells colored by their loadings. **(m)** Cell type distributions of most weighted component 8 receiver cells (top 1%).

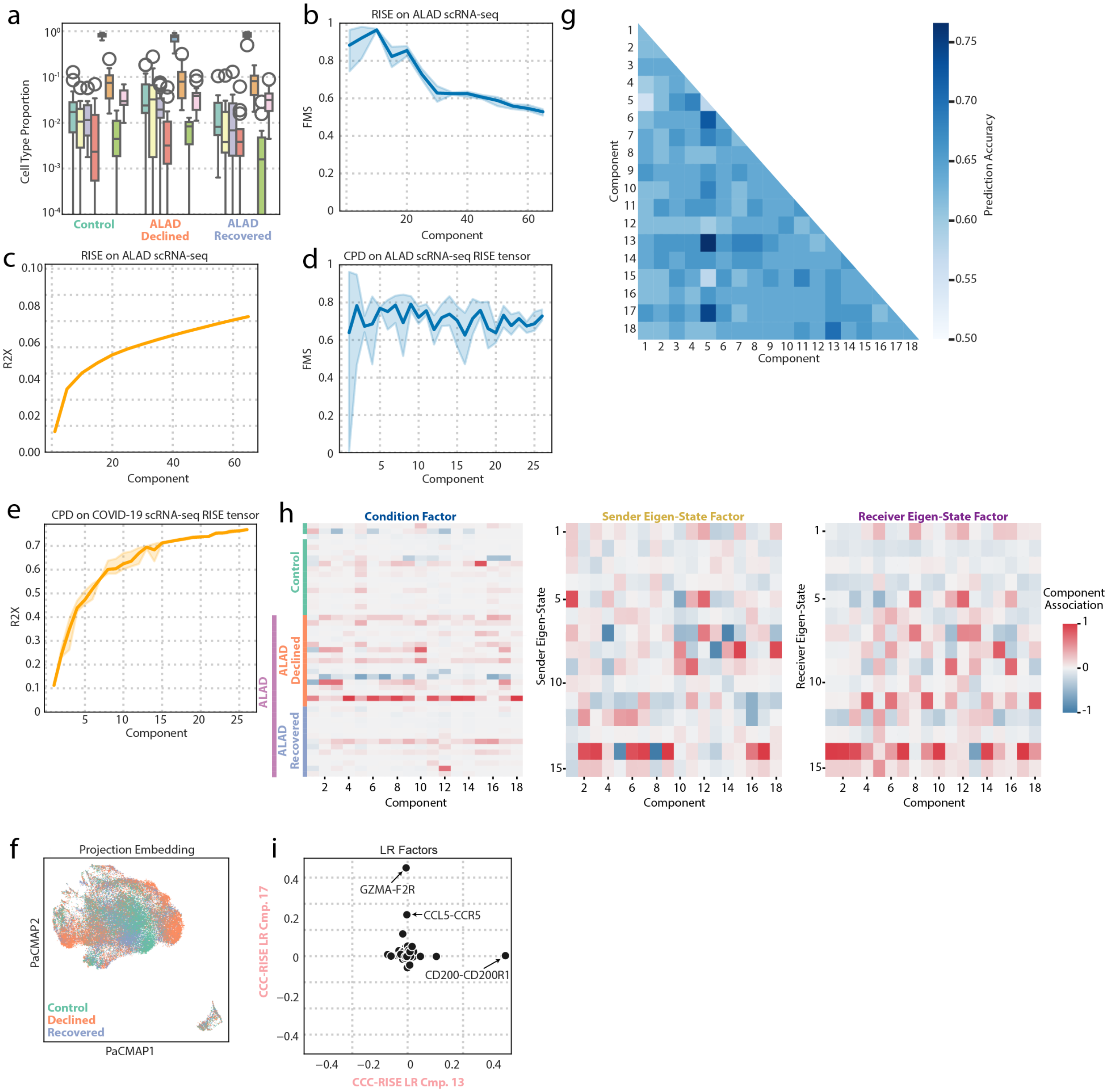


**Figure S5. Validation of the CCC-RISE model and characterization of the ALAD patient cohort.** **(a)** Cell type proportion (CD4 T cells, CD8 T cells, dendritic cells, epithelial cells, macrophages, monocytes, NK cels, and proliferating cells) across the patient categories. **(b)** FMS when comparing the RISE gene and condition factors of the full scRNA-seq COVID-19 dataset to a bootstrapped dataset at various ranks. **(c, e)** Fraction of the variance explained by (g) RISE and (i) CPD for each number of components. **(d)** FMS when comparing the CPD condition and sender/receiver eigen-state factors of the 4D communication eigen-state tensor (15-component RISE model) to a bootstrapped tensor at various ranks. **(f)** PaCMAP of the RISE projections with cells colored by their cell-type annotations. **(g)** ALAD prediction accuracy of an unpenalized logistic regression model trained on pairs of components. **(h)** The condition, sender eigen-state, and receiver eigen-state factors. **(i)** Correlation between CCC-RISE component 13 and component 17 LR pairs with the most highly weighted LR pairs annotated.

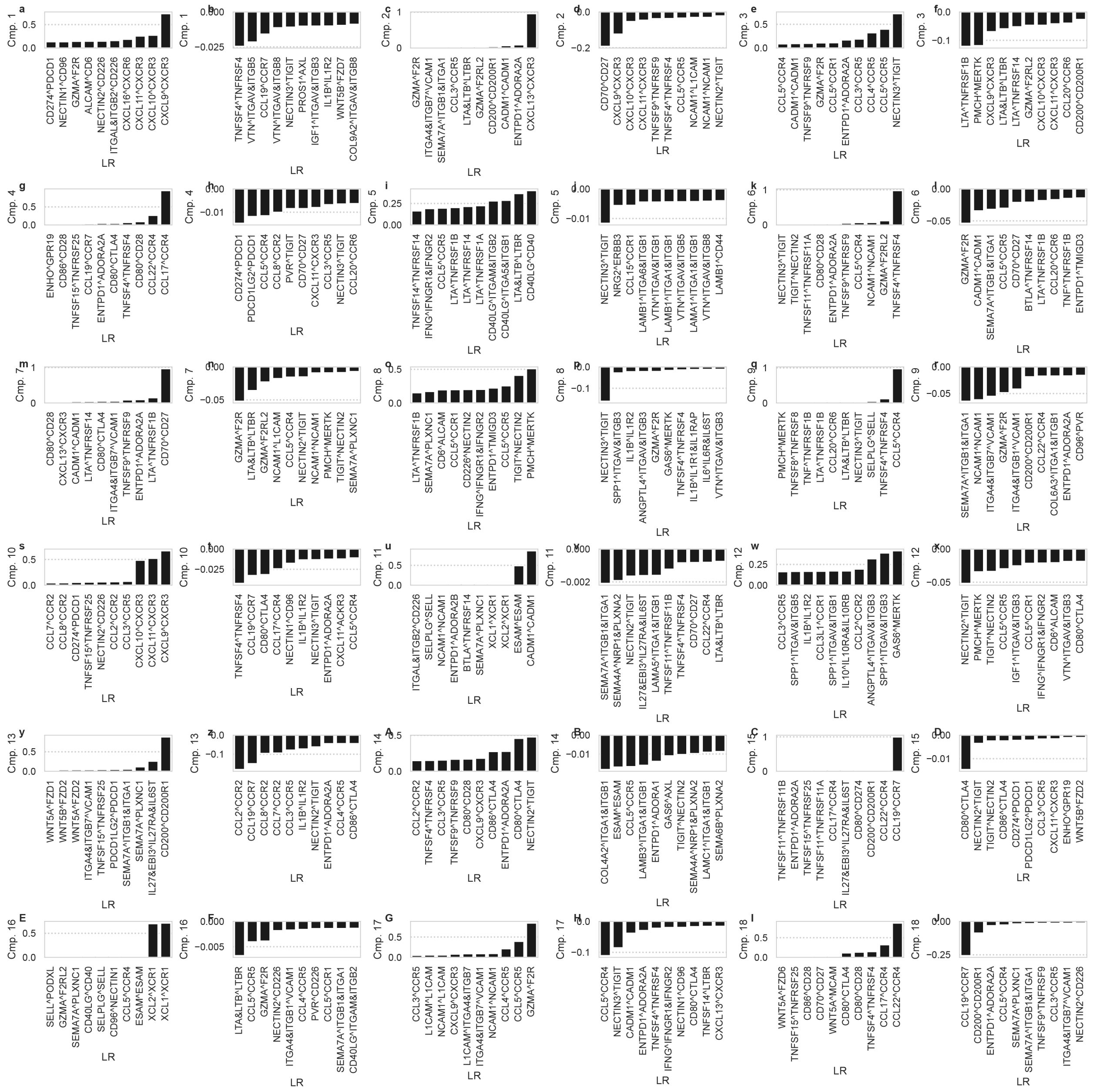

**Figure S6. CCC-RISE components associate each pattern to specific LR pairs for the scRNA-seq ALAD dataset. (a, c, e, g, i, k, m, o, q, s, u, w, y, A, C, E, G, I)** The top 10 most positively weighted LR pairs identified by CCC-RISE by a 35-component RISE and 8-component CPD model. **(b, d, f, h, j, l, n, p, r, t, v, x, z, B, D, F, H, J)** The top 10 most negatively weighted LR pairs identified by CCC-RISE by a 15-component RISE and 18-component CPD model.

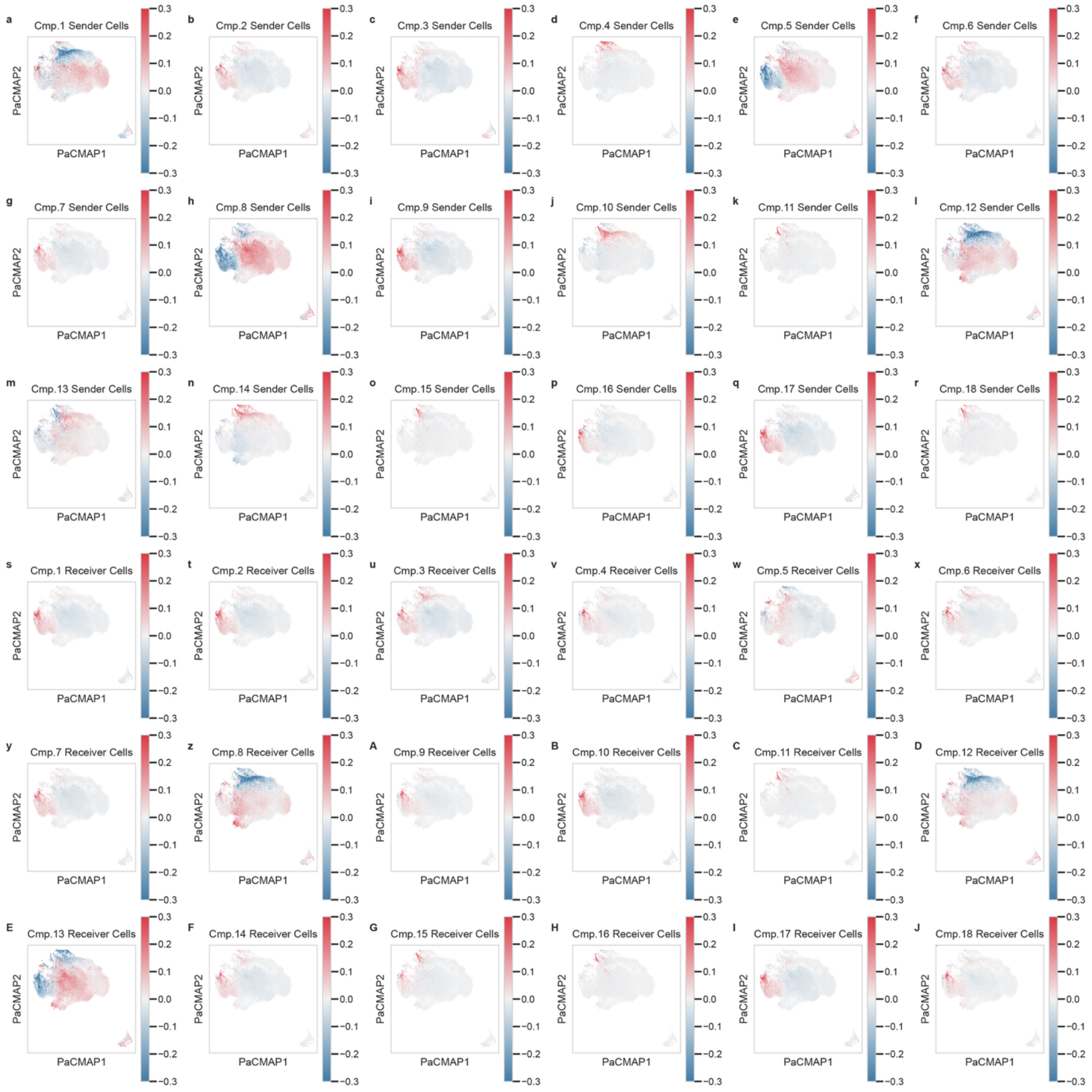


**Figure S7. CCC-RISE components associate each pattern to specific sender and receiver cells for the scRNA-seq ALAD dataset for a 15-component RISE and 18-component CPD model. (a-J)** PaCMAP of the RISE projections, with (a-r) sender cells and (s-J) receiver cells colored by the weightings for the indicated component.

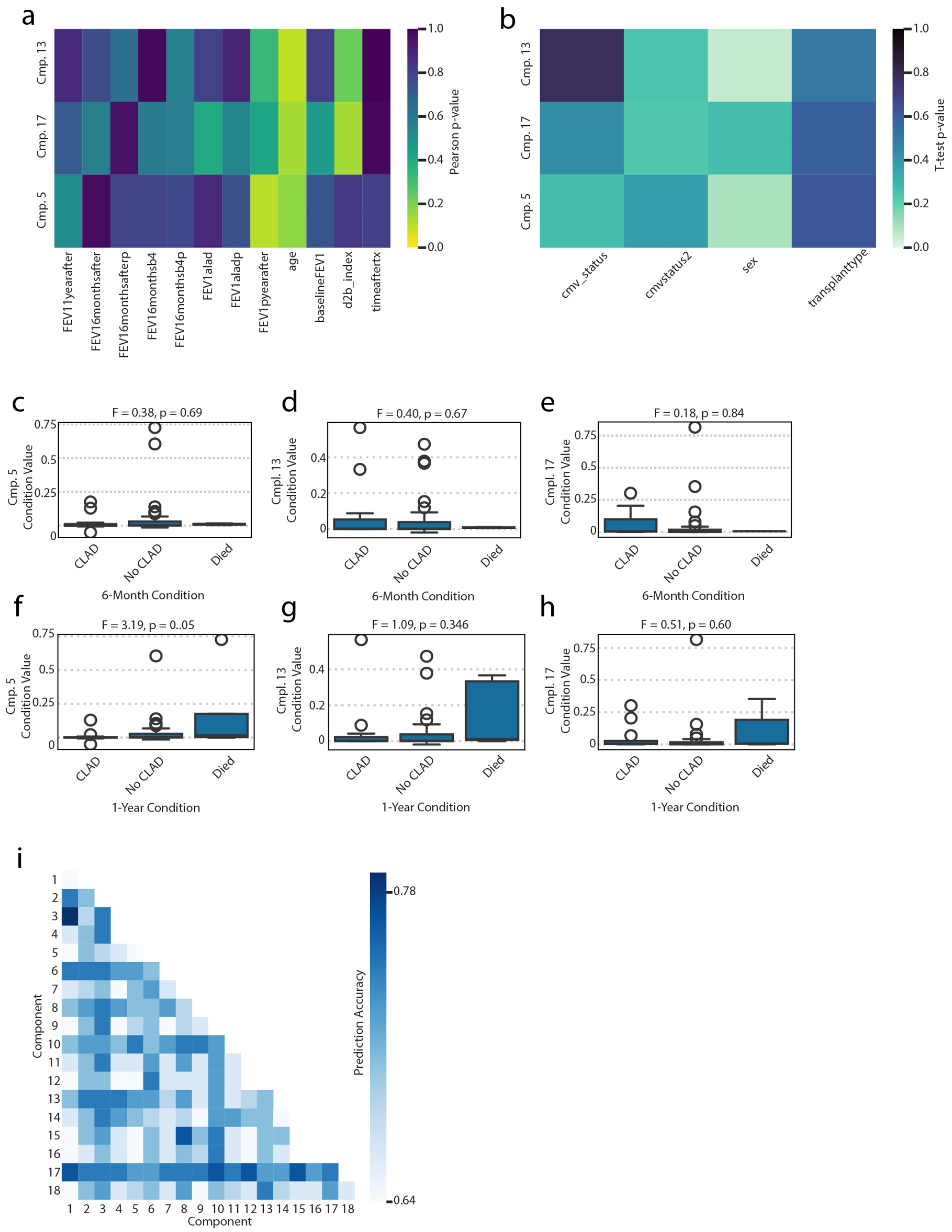


**Figure S8. Association of specific CCC-RISE components with clinical metadata and outcomes. (a)** Pearson correlation p-value between condition loadings for components 5, 13, and 7 and clinical variables. **(b)** T-test p-value between condition loadings for components 5, 13, and 17 stratified by categorical clinical information. **(c, f)** Component 5 condition weights stratified by (c) 6-month CLAD status and (f) 1-year CLAD status. **(d, g)** Component 13 condition weights stratified by (d) 6-month CLAD status and (g) 1-year CLAD status. **(e, h)** Component 17 condition weights stratified by (e) 6-month CLAD status and (h) 1-year CLAD status. **(i)** Stable prediction accuracy of an unpenalized logistic regression model trained on pairs of components.

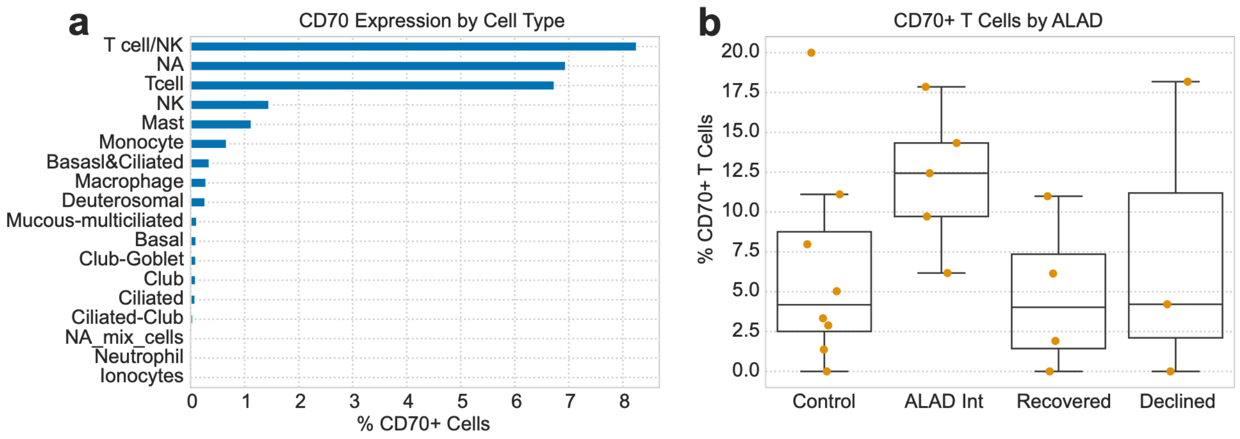


**Figure S9. A subset of CD70-positive cells is a reproducible feature of ALAD. (a)** The percentage of CD70-positive cells within the airway brush samples dataset, subset by cell type. **(b)** The percent of T cell CD70 positivity within each sample, subset by clinical context.
